# Universal statistics of hippocampal place fields across species and dimensionalities

**DOI:** 10.1101/2024.06.11.597569

**Authors:** Nischal Mainali, Rava Azeredo da Silveira, Yoram Burak

## Abstract

Hippocampal place cells form a spatial map by selectively firing at specific locations in an animal’s environment^1^. Until recently the hippocampus appeared to implement a simple coding scheme for position, in which each neuron is assigned to a single region of space in which it is active^1^. Recently, new experiments revealed that the tuning of hippocampal neurons to space is much less stereotyped than previously thought: in large environments, place cells are active in multiple locations and their fields vary in shape and size across locations, with distributions that differ substantially in different experiments^2–7^. It is unknown whether these seemingly diverse observations can be explained in a unified manner, and whether the heterogeneous statistics can reveal the mechanisms that determine the tuning of neural activity to position. Here we show that a surprisingly simple mathematical model, in which firing fields are generated by thresholding a realization of a random Gaussian process, explains the statistical properties of neural activity in quantitative detail, in bats and rodents, and in one-, two-, and three-dimensional environments of varying sizes. The model captures the statistics of field arrangements, and further yields quantitative predictions on the statistics of field shapes and topologies, which we verify. Thus, the seemingly diverse statistics arise from mathematical principles that are common to different species and behavioral conditions. The underlying Gaussian statistics are compatible with a picture in which the synaptic connections between place cells and their inputs are random and highly unstructured.

## Main

A central goal of systems neuroscience is to characterize how neural systems represent information about the world, or about the brain’s internal state. For several decades, much of the thinking about neural population codes was motivated by reports on neurons with highly stereotyped tuning functions. Neurons were often observed to have a smooth, typically unimodal tuning to the encoded variable, centered around preferred stimuli that vary across the neural population^8,9^. Examples include the radially symmetric receptive fields of retinal ganglion cells^10^, receptive fields of simple cells in the primary visual cortex^11,12^, cosine tuning to movement direction in the motor cortex^13^, unimodal fields of head direction cells^14^, and the fields of hippocampal place cells in small environments^1^.

Experiments in the past decade, however, have uncovered neural response functions that are much less stereotyped and regular than observed previously^2,3,15–21^. These recent findings were enabled by high-throughput recording techniques which substantially reduced bias in the selection of cells for analysis, as well as new technologies that enabled the monitoring of neural activity in freely behaving animals, under reduced behavioral constraints^6,22–24^.

Some of the most striking examples of irregular neural responses were recently identified in area CA1 of the hippocampus^2–7^. The classical view of spatial coding by place cells in this area has been that they are active in a single, compact region of space, and exhibit a stereotyped bell-shaped tuning to position^1,25^. Several recent experiments in bats and rodents have revealed, however, that this picture breaks down in large environments. Place cells typically fire in multiple locations and, furthermore, the multiple firing fields of individual cells, as well as those of the whole population, vary in size and in their shape, which can deviate substantially from the classical bell-shaped form^2–7^. It remains unknown, however, whether the irregular statistics of place fields in bats and rodents can be described in terms of common mathematical principles, and whether these statistics yield insights on the synaptic architecture that underlies spatial coding in hippocampal area CA1.

Here, we report that a surprisingly simple generative model accounts for highly detailed features of place field statistics. We model place fields as derived from a realization of a random Gaussian process over the spatial coordinates: a realization of the process is sampled for each cell and is then thresholded and rectified, resulting in multiple, heterogeneous fields (Fig 1A). A Gaussian process is a random function, whose values at any discrete set of positions are jointly normal^26^. The statistics of a Gaussian process are uniquely determined by its mean, which we set to zero without loss of generality as its choice is redundant with the choice of the threshold, and by the spatial correlation function, which we assume to be translationally invariant.

**Fig. 1.**
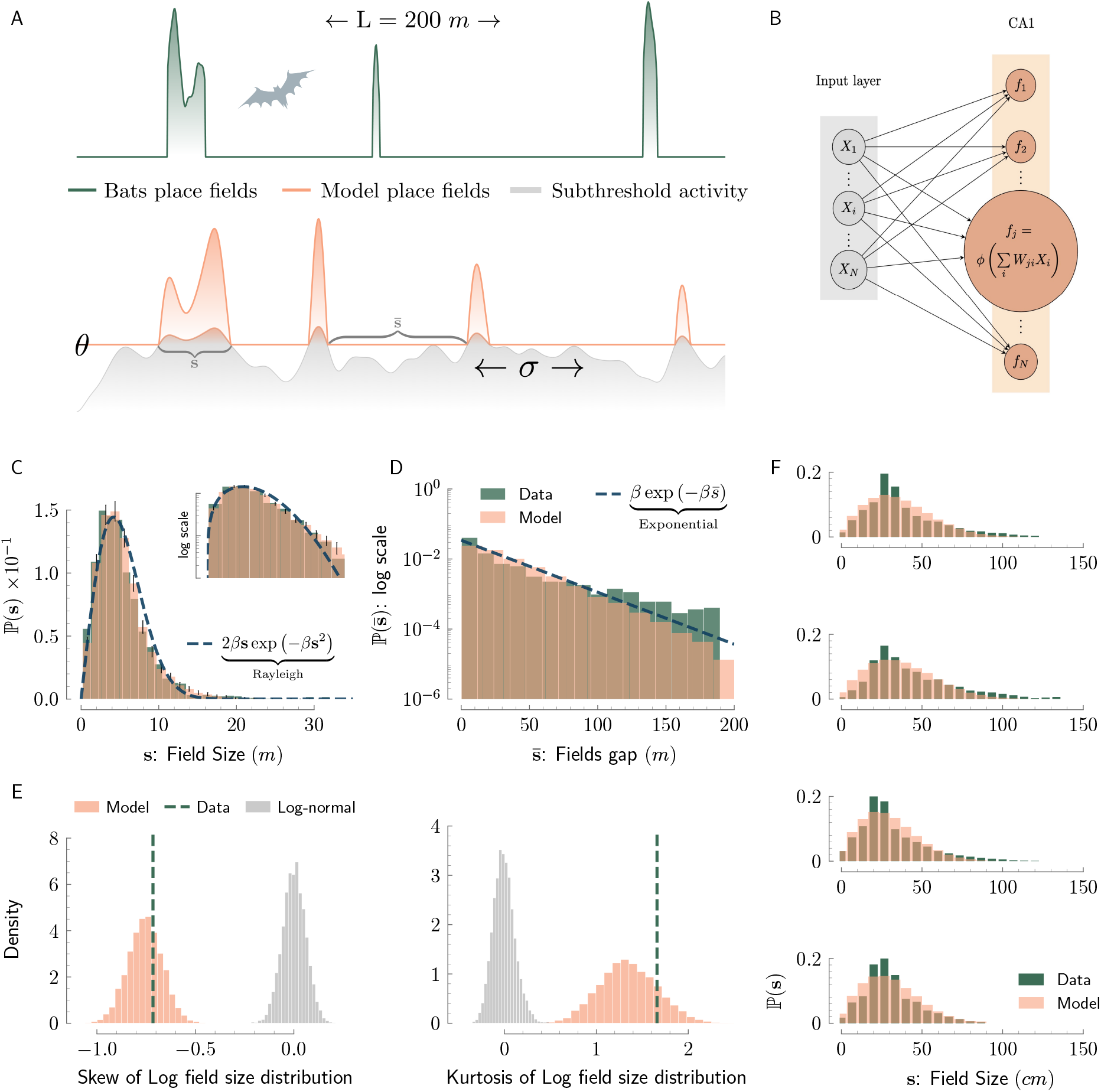
Field arrangements of 1d place fields are explained by the thresholded Gaussian process model. **a**, Top: example of place fields measured in bats flying in a 1d 200 m-long tunnel^6^. Bottom: firing rates in the model are generated by thresholding and rectifying a realization of a random Gaussian process (gray). In all figures, green represents experimental data, and orange represents the model. **b**, A simple neural architecture which gives rise to place fields that follow the thresholded Gaussian process model. Presynaptic inputs from many spatially selective neurons are summed with random synaptic weights, to create the input to each CA1 place cell, followed by rectification. **c**, Field size distribution of bats in the 1d tunnel compared against the field size distribution predicted by the model. Dotted line: approximate analytical prediction of the field size distribution, valid for high thresholds (Rayleigh distribution). Orange bars: precise predictions of the model, obtained from simulations (error bars: standard deviation across simulations, see methods). Here and in all subsequent bar-plots, the overlap between green bars (experiment) and orange bars (model) is represented in brown. Inset: same data in semi-logarithmic scale. **d**, Distribution of consecutive field gaps in the experiment compared against the distribution predicted by the model. Dashed line: analytical prediction. **e**, Skew (left) and kurtosis (right) of the log-field-size distribution, obtained from multiple simulations of the model and the best-fit heuristic log-normal distribution (gray). Each simulation is matched to the size of the experimental data set. For the log-normal distribution, the empirical skew and kurtosis are expected to be close to zero. Dashed line: skew and kurtosis of the experimental distribution. **f**, Rayleigh distribution compared against place field size distribution in four different mice, running in a 40 m-long virtual track^5^.

Two lines of reasoning have led us to examine the above model. First, a Gaussian process is the random process that maximizes entropy under constraints on the mean and the spatial covariance function. Hence, our modelling choice invokes minimal assumptions other than a specification of a place cell’s underlying input correlation structure^27^. Second, Gaussian statistics arise when many independent random variables are summed. The summed presynaptic input into area CA1 from area CA3 and entorhinal cortex is thus expected to be approximately Gaussian if synaptic weights are predominantly random (Fig. 1; see supplementary information A)^28^. We further discuss these motivations in the Discussion.

### Field arrangements

Place field locations in the model are identified as the regions in space in which the Gaussian process exceeds a given threshold. Extensive results exist on the statistics of these regions in the mathematical literature, where they are called *excursion sets*^26^. A recent result of central importance to this work is that threshold crossing statistics assume a universal form when the spatial correlation function of the process obeys mild smoothness requirements, and the threshold is sufficiently high^29^. Threshold-crossing statistics are then insensitive to the detailed structure of the correlation function, and depend only on two scalar parameters: the normalized threshold, *θ*, defined as the ratio between the threshold and the standard deviation of the Gaussian process (Eq. S2) and a correlation length, *σ*, which is derived from the spatial correlation function (Eq. S3).

Specifically, the mean density of fields and the mean field size can both be expressed in terms of *σ* and *θ* using the Kac-Rice formula^30^ for the threshold crossings of a Gaussian process (Eqs. S20 and S21). Furthermore, the full distribution of field sizes acquires a universal form in the limit of large *θ*, expressed as

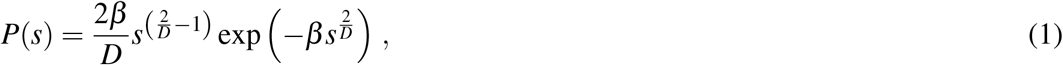

where *D* is the dimensionality of the space over which the Gaussian process is defined, *s* is the field size (length in one dimension, area in two dimensions, and volume in three dimensions; 1d, 2d, and 3d hereafter), and the single parameter *β* can be expressed in terms of *σ* and *θ* (Eq. S25; see supplementary information B). In the 1d case (*D* = 1), this expression takes the form of a Rayleigh distribution (Fig. 1C). Similarly, field gap statistics assume a universal form in the high-threshold limit: threshold crossings are spatially uncorrelated and, consequently, gap sizes follow an exponential distribution, with a mean that can be expressed in terms of *σ* and *θ* (see methods). Thus, according to the model, the empirical distributions of CA1 field sizes and gaps between fields should be jointly explained using the two parameters, *σ* and *θ* .

We began by examining place fields recently recorded in bats flying in a 200 m-long tunnel, where the highly stereotyped trajectory of the animals along the 1d track allowed for the gathering of comprehensive statistics on the spatial arrangement of place fields^6^. We determined the two parameters *σ* and *θ* by matching the means of the field sizes and gaps to the model in the data from the 200 m-long tunnel. This fitting procedure, based only on the means, was sufficient to capture the full distributions of field sizes and gaps (Figs. 1C and 1D). This outcome was particularly striking for the field sizes, whose distribution had a distinctive, highly asymmetric form.

Due to its highly asymmetric structure (Fig. 1C), the field size distribution was previously fitted heuristically to a log-normal distribution, yet this approach lacked a principled rationale. By contrast, in our model, the asymmetry is explained by the statistical dependencies between locations of adjacent threshold crossings of the underlying random Gaussian process^31^. The empirical field size distribution was more likely to arise from the Gaussian threshold-crossing model than the best-fit log-normal distribution, even though the model had one fewer degree of freedom (assessed via likelihood and parameter-aware likelihood based model performance metrics: ΔLLR *<* 0, ΔBIC *<* 0, ΔAIC *<* 0; see methods). Furthermore, skew and kurtosis of the log-field-size distribution were in agreement with the threshold crossing model, but not with the log-normal distribution, according to which they should be close to zero (Fig. 1E; see methods). Thus, the model captured subtle features of the distribution, beyond its first and second order moments. As expected in the high-threshold limit, the distribution of gap sizes followed an exponential distribution (Fig. 1D).

We subsequently tested the model’s ability to explain the statistics of place fields measured in rodents running in long 1d tracks. Field size distributions extracted from experiments in which mice ran in a 40 m-long 1d virtual track^5^ were in qualitative agreement with the Rayleigh distribution (Fig. 1F). We also analyzed the distribution of field sizes from an experiment in which rats exlored a 48 m-long 1d maze^3^. The distribution of field sizes was in quantitative agreement with the distribution predicted by the model, with matching skew and kurtosis (Extended Data Fig. 1).

### Place fields in two and three dimensions

Experimental data on place fields in rats navigating large 2d environments^7^ and bats in 3d rooms^4^ reveal place cells with multiple, heterogeneous place fields. Intriguingly, though, the experiments uncover field size distributions that are distinct in one, two, and three dimensions. Building on our initial findings in 1d environments, we examined the extension of the model to 2d and 3d environments, where place fields are determined by excursion sets of a Gaussian process over the higher-dimensional space. As in the 1d case, the model generated multiple, heterogeneous fields, which were qualitatively similar to the ones observed experimentally (Figs. 2A-B and 2F-G).

**Fig. 2.**
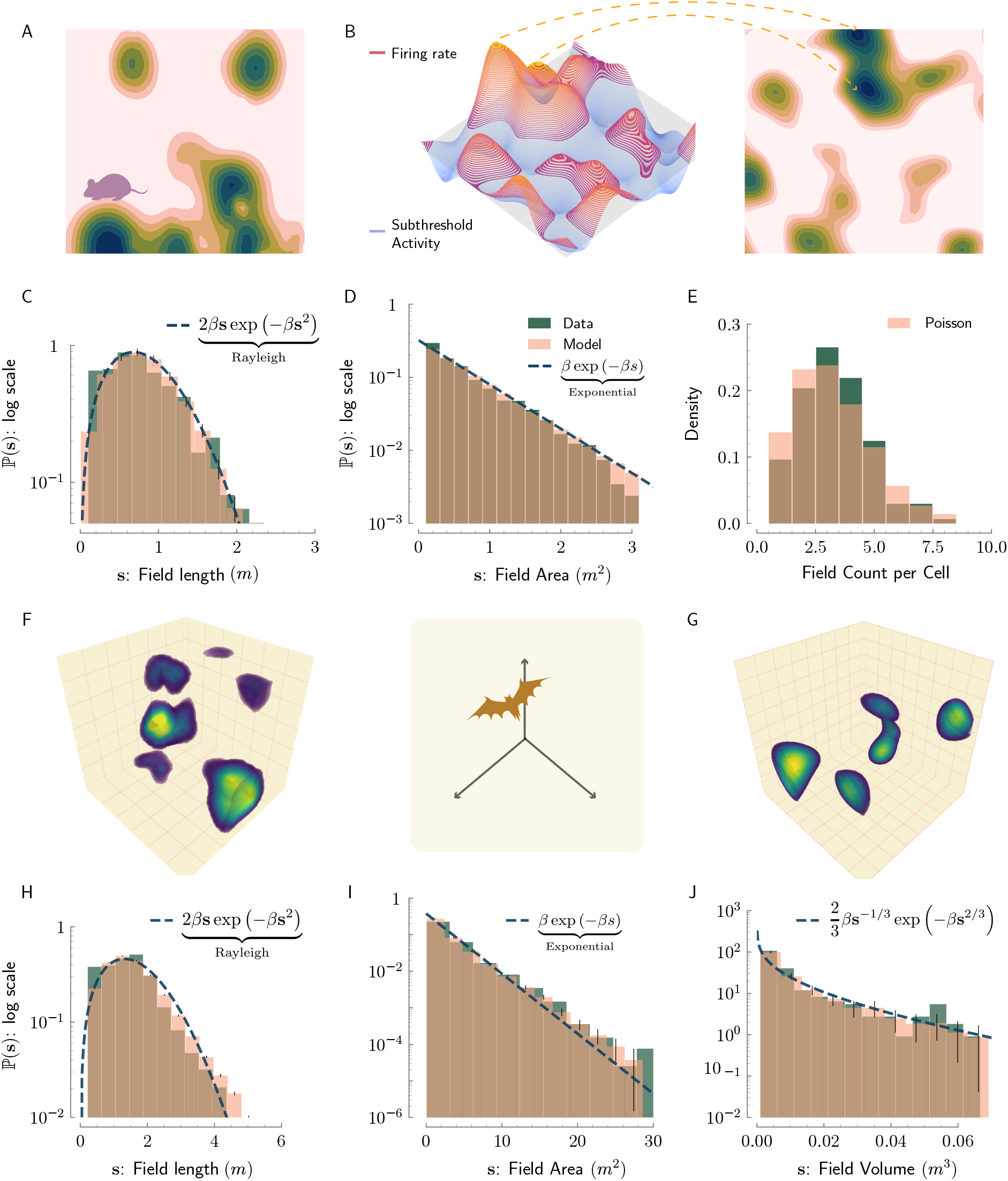
Field arrangements of place fields in 2d and 3d spaces are explained by the thresholded Gaussian process model. **a-e**, Model fits to 2d data from rats. **a**, Sample spatial firing pattern of a CA1 cell, recorded in a rat navigating in a 18.6 m^2^ environment^7^. **b**, Model: Random Gaussian process in 2d (left) is thresholded to generate firing fields, yielding the cell’s spatial firing pattern (right). In all the panels below that show distributions, the vertical scale is logarithmic and the color scheme for comparison between the model and the experiment is as in Fig. 1C, 1D. **c**, Distribution of field lengths in 1D slices of 2d place fields (20 sample cells from^7^). **d**, Distribution of place field areas from all cells in the experiment (experimental histogram adapted from^7^). **e**, Distribution of field count per cell across the whole population of recorded cells (experimental histogram adpated from^7^). **f-j**, Model fits to 3d data from bats. **f**, Sample 3D firing pattern recorded in a bat CA1 cell during flight in a 5.8 × 4.6 × 2.7 m^3^ room^4^. **g**, Visualization of 3D receptive fields generated by the model. **h**, Distribution of field lengths in 1D slices of bat 3D place fields (analysis performed on all recorded cells in^4^). **i**, Distribution of field areas in 2D slices of the bat 3D place fields. **j**, Distribution of 3D field volumes in bats. As the dimension increases, the distribution becomes more concentrated around zero, and develops a longer tail.

Moreover, 1d slices through a multidimensional Gaussian process are themselves Gaussian processes. Hence, field sizes in 1d slices through multidimensional firing fields are expected to follow the same field size distribution as in the 1d case. To test this prediction, we examined 1d slices through 2d place fields measured in rats foraging in a 18.6 m^2^ arena (data from 20 cells recorded in^7^), and found that they were well fit by the Rayleigh distribution (Fig. 2C). We further compared the model against a histogram of field areas from all cells (as reported in^7^). The statistics precisely followed an exponential distribution as predicted by the model (Eq. 1, D = 2; Fig. 2D). Finally, field counts were well fit by a Poisson distribution, as expected in the high-threshold regime in which field locations are independent (Fig. 2E).

Next, we analyzed 3d place fields in bats navigating a large room of size 5.8 × 4.6 × 2.7 m^34^. The model predicts that 2d slices through the 3d place fields should exhibit the same statistics as those of fields in 2d environments, since these slices are also realizations of a thresholded Gaussian process in two dimensions. The predictions for both 1d and 2d slices were verified by the data (Figs. 2H-I), along with the distribution of the 3d field volumes (Fig. 2J), which followed the prediction of Eq. 1 with *D* = 3. In an alternative model, in which field volumes and peak firing rates were exactly matched to the empirical data, without the underlying Gaussian statistics, distributions of 1d and 2d slice sizes did not match the data (Extended Data Fig. 2).

In summary, the statistics of field arrangements in 1d, 2d, and 3d were explained in both bats and rodents by the threshold crossings of Gaussian processes. In particular, the model explained the different field size statistics that were observed across experiments in different dimensionalities (Eq. 1 and Figs. 1C, 2D, and 2J). The generality of these results suggests that common mechanisms, shared across species and dimensionalities, underlie the structure of the hippocampal code for space.

### Statistics of field shapes

The statistics of field sizes and gaps only reflect the statistics of field boundaries which, in the model, correspond to threshold crossings. The model also makes quantitative predictions on the statistics of field shapes, defined as the firing rate’s dependence on position within place fields (Fig. 3A). These statistics are determined by the properites of the Gaussian process in the threshold-crossing segments. First, according to the model, the firing rate within a given place field may exhibit multiple local maxima, unlike in the classical picture of bell-shaped fields. The predicted distribution of the number of local maxima across fields was in close agreement with the empirical distribution of local maxima per field in bats flying in the 1d tunnel (Fig. 3B). Second, the model predicts a positive correlation between the size of a field and its peak firing rate, with an approximate power-law relation between these two quantities. The experimental data confirmed this prediction, both in bats flying in the 1d tunnel (Fig. 3C) and bats flying in the 3d environment (Fig. 3D). In both cases, the model not only predicted the qualitative relation between field size and peak firing rate, but it also accurately predicted the power law coefficient (Figs. 3C and 3D). Third, in the model, the slope of the firing rate at the boundaries of 1d place fields follows a Rayleigh distribution^32^. This prediction was verified in the data (Fig. 3E). Fourth, in bats flying in the 3d environment, the mean and Gaussian curvatures of iso-surfaces formed by the boundary of 3d place fields were distributed in agreement with the predictions of the model (Extended Data Fig. 3). All the above predictions of the model were verified using the values of *σ* and *θ* that were previously inferred based only on field arrangements, independent of field shapes.

**Fig. 3.**
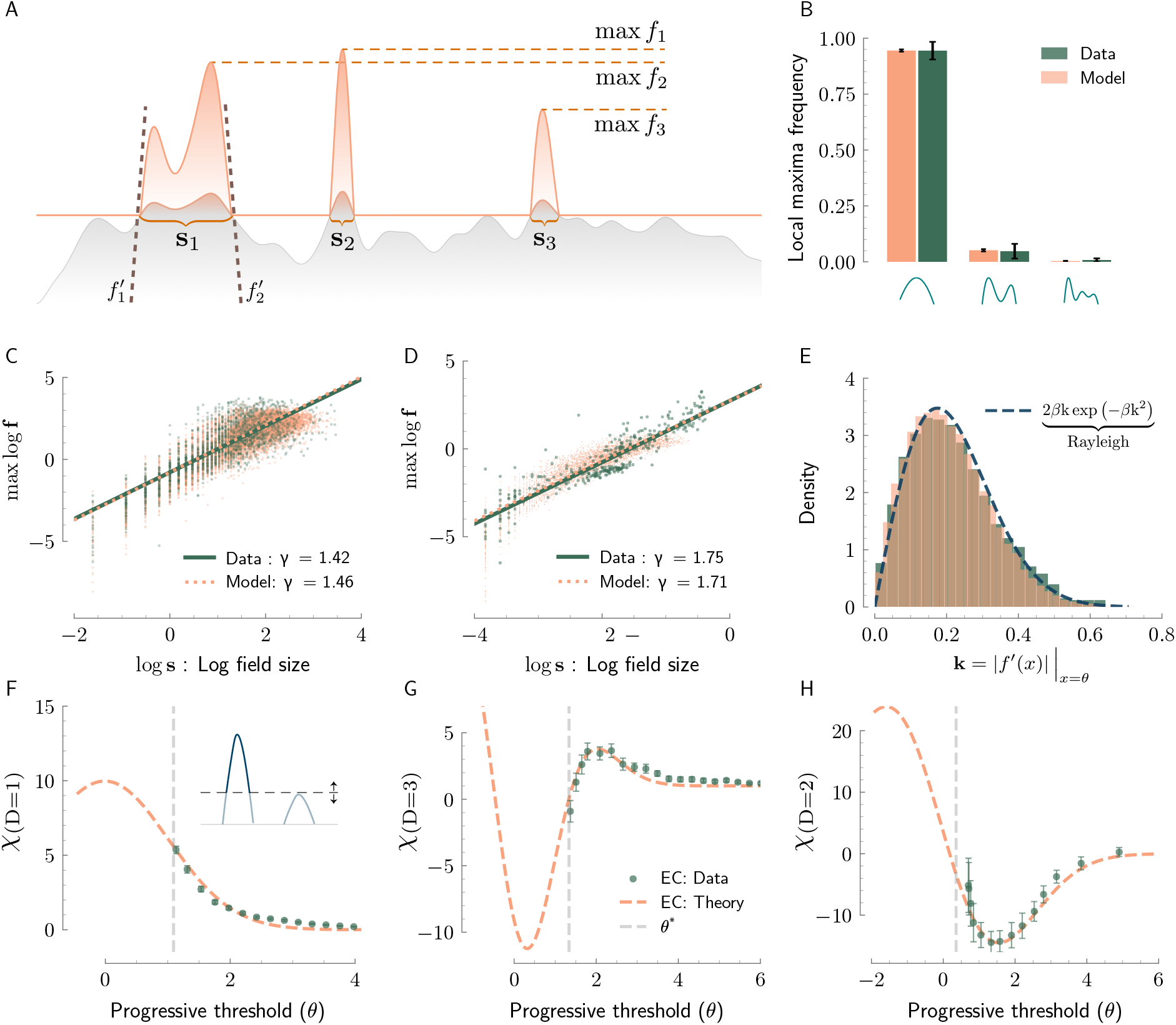
Field shape related statistics of place fields are predicted by the thresholded Gaussian process model across dimensions and species. **a**, Visualization of various field shape related measurement in case of 1d fields. **b**, Distribution of the number of peaks in the place fields of bats in the 200 m-long tunnel^6^ compared to the model. Error bars: standard deviation of the mean (model) and standard deviation across a range of smoothing parameters (data, see methods). **c**, Joint distribution of log receptive field size (in m) and log maximal firing rate (in Hz) obtained from the model (orange) and from experimentally measured bat place fields in the 200 m-long tunnel (green). Lines: power law relationship between the two quantities, in the model and experiment. The coefficient *γ* is the slope of the best linear fit to the distribution and represents the power law coefficient. The y-intercept of the fit is matched between data and model to infer the linear gain factor. **d**, Same as in A, for place fields recorded in bats flying in the 5.8 × 4.6 × 2.7 m^3^ room^4^. Field size is the cubic root of the place field volume (in m). **e**, Absolute value of field slopes at their boundaries follow a Rayeligh distribution. **f**, Euler characteristic (*χ*) as a function of extra thresholding of the place fields of bats in the 200 m-long tunnel (Dashed line: theoretical prediction, Dots: data). Inset illustrates how adding an extra threshold (horizontal dashed line vs. solid line) can affect the number of fields. **g**, Similar plot as D for bats in 3D environment^4^. **h**, Same as D,E in the 18.6m^2^ 2d environment (using 20 sample cells from rats^7^).

We next examined whether the model captures topological properties of the place fields, using recent results on the Euler Characteristic (EC) of excursion sets of Gaussian processes. The EC of the thresholded Gaussian process is a function of the number of connected components and holes of various dimensions (see methods), and has a universal dependence on the threshold which is specific to Gaussian processes^26^. An analytical expression has been obtained for this quantity in terms of *θ* and *σ*. (This results has also been used to test for Gaussianity in the spatial structure of the cosmic background microwave radiation, as well as other natural processes^33–36^.)

In the 1d case, the EC is simply equal to the number of connected components. The theory provides an exact expression for the EC curve (Eq. S7), whose only dependence on the correlation function is through the correlation length *σ*. We tested this prediction for place fields by placing a varying threshold on the firing rates measured in bats flying in the 200 m long tunnel, followed by evaluation of the EC of the rectified fields. The resulting EC curve (Fig. 3F) was in excellent agreement with the analytical prediction using the previously obtained value of *σ* .

In dimensions higher than one the EC depends not only on the number of connected components but also on the number of holes of various dimensions within the fields. This results in a non-monotonic dependence of the EC curve on the threshold, due to the emergence and disappearance of holes. Here, too, the theory provides an exact analytical expression (Eq. S9) for the expected EC curve. We tested this prediction on all measured firing fields in bats flying in 3d^4^. The empirical EC curve was non-monotonic as predicted, and precisely followed the analytical expression (Fig. 3G) using the previously obtained value of *σ*. This agreement is specific to the Gaussian threshold crossing model, and was not reproduced by an alternative model with matching joint statistics of field volumes and peak rates (Extended Data Fig. 2C).

We also tested the EC curve prediction in 2d, on a data set of 20 cells from rats recorded in the 2d megaspace^7^, and obtained excellent agreement with the analytical prediction (Eq. S8; Fig. 3H).

### Parameter variation across experiments

Our analyses of data from 1d and 2d environments of varying sizes allow us to identify qualitative trends across datasets, as a function of the environment size (Figs. 4A and 4B). We observe that the correlation length, *σ*, increases sublinearly with the size of the environment, and the normalized threshold, *θ*, decreases with the size of the environment (Fig. 4B). Together, these trends induce an increase in the average number of fields with the size of the environment, consistent with the observation of single firing fields per cell in small environments, and a sublinear increase in their sizes (Extended Data Fig. 4).

**Fig. 4.**
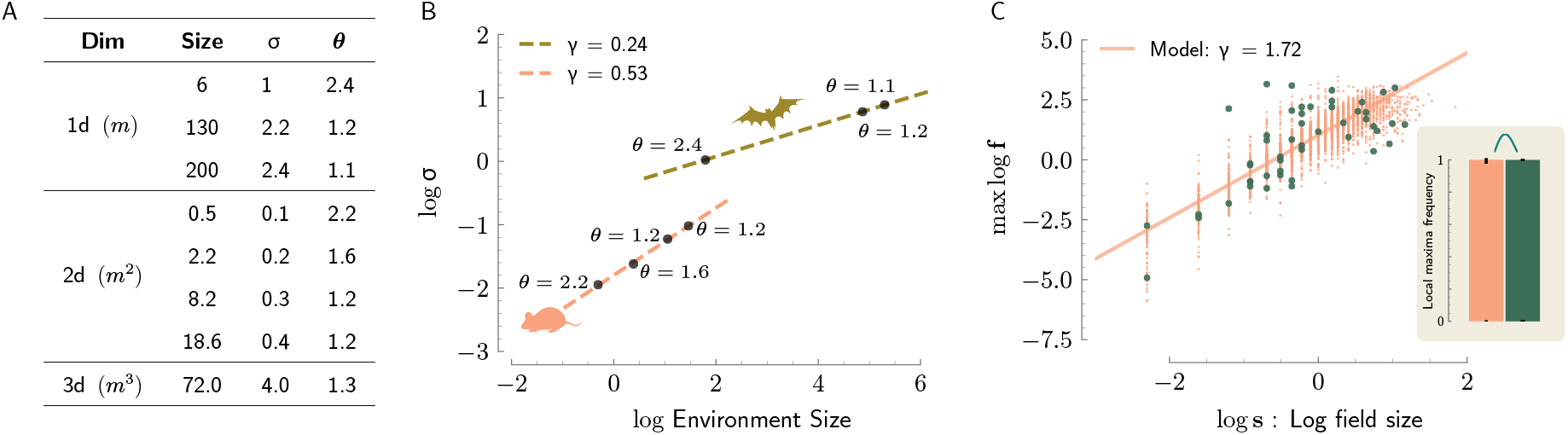
Parameter trends across experiments explain differences in heterogeneity. **a**, Summary of parameters for the various experiments in spaces of varying dimension: 1d, bats flying in a tunnel; 2d, rats navigating a large open arena; and 3d, bats flying in a room. **b**, Linear scaling of log *σ* with the logarithm of the linear environment size in 1d (bats) and 2d (rats). **c**, Joint distribution of log receptive field size, *s* (in m) and log maximal firing rate, *f* (in Hz), plotted as in Fig. 3C, but in the 6 m-long tunnel. In comparison with the 200 m-long tunnel (Fig. 3C), the exponent characterizing the relationship between field size and maximal firing rate is closer to 2, indicating that field shapes in the 6 m-tunnel are more stereotyped than in the 200 m-long tunnel, as expected due to the higher threshold. Inset: fields with multiple peaks are practically absent, as predicted (compare with Fig. 3B). In panels B and C, *γ* is the slope of the best linear fit to the data in logarithmic scales.

While the two parameters jointly affect the field counts and the field sizes, the normalized threshold also influences the heterogeneity in field shapes (Extended Data Fig. 5A). With increasing *θ*, the fraction of multi-peaked fields decreases and the exponent of the power-law relationship between the maximum firing rate and the field size approaches 2, as expected for parabolic fields^26^ (Extended Data Figs. 5B and 5C). Thus, the model predicts greater stereotypy of field shapes with increasing normalized threshold. As a result, and following the trend seen in the data (Figs. 4A and 4B), field shapes are predicted to be more stereotyped in smaller environments than in larger ones (Extended Data Figs. 5A-C). We tested this prediction on place fields measured in bats flying in a short, 6 m-long tunnel^6^. As predicted, field shapes were more stereotyped than in the 200 m-long tunnel, with statistics that quantitatively agreed with the model (Figs. 4C and Extended Data Fig. 5D). In summary, the model accounts for the different characteristics of CA1 spatial selectivity across environmental scales, ranging from early experiments in small environments, where place cells typically had a single, bell-shaped firing field^1^ to more recent experiments in large environments, where place cells had multiple firing fields with heterogeneous shapes^3,5–7^.

Each one of the experimental data sets is well explained by a single choice of the values of *σ* and *θ*, yet some variation in these parameters across cells and anatomical regions is to be expected. For the length scale *σ*, our choice of a single value is motivated by the fact that recordings were tightly localized anatomically in dorsal CA1^4,6,7,37^. In the data from bats flying in the 1d tunnel, where comprehensive measurements allow for fine exploration of the statistics, the number of place fields expressed by individual neurons, as well as their sizes, were observed to be correlated in the two flying directions, despite remapping of the fields^6^. We hypothesized that this correlation arises from variability in the threshold, *θ*. In agreement with this hypothesis, an extended model with a distribution of thresholds accounted for the empirical correlations, while remaining compatible with the distributions of field sizes and gaps obtained from the basic model (Extended Data Fig. 6).

## Discussion

The classical view of spatial coding in the hippocampus has been challenged by the recent discoveries of heterogeneous and distributed CA1 response patterns in large environments. It was unknown whether a unified mathematical framework can encompass the diverse statistics of these irregular responses. Here, we identified such a framework, in which place fields are obtained from the threshold crossings of a spatially fluctuating random process. The underlying randomness of the Gaussian process enabled the model to explain in quantitative detail the variability in the size of place fields and their spatial arrangement, as well as a range of geometric and topological properties associated with the heterogeneous field shapes. Indeed, when constrained only on the mean field size and count, the model precisely captured the statistics of shape variability, which in principle could have had an independent origin. This surprising result suggests that the different degrees of heterogeneity, observed across experiments, arise as a byproduct of mechanisms that regulate the field frequency and size. With only two parameters, the model accounts for the seemingly diverse statistics observed in rodents and bats, and in environments of varying sizes and dimensionalities. Therefore, the model allows for direct comparison between the statistics observed in the different experiments, and suggests that they arise from common underlying principles. A Gaussian process is the random process that maximizes entropy under constraints on the mean and the spatial covariance function. Hence, the model invokes minimal assumptions other than a specification of a place cell’s underlying input correlation structure^27,38^. At the same time, random Gaussian processes can generate a highly efficient coding scheme, with capacity that increases exponentially with the number of neurons^28^. It will be interesting to examine theoretically whether it is possible to obtain a coherent understanding of the trends seen in the parameters across the different geometries (Fig. 4B) in light of a theory of efficient coding^28^.

The success of the parsimonous model introduced here invites an exploration into its possible mechanistic origin. The apparent stereotypy of place fields in small environments has motivated a view of hippocampal spatial selectivity as arising from highly organized synaptic connectivity^39–41^. However, the irregular CA1 firing patterns in large environments are inconsistent with this view. Our results indicate that CA1 place fields, in small and large environments alike, are compatible with predominantly random synaptic connections into CA1. This is because random projections from a sufficiently large number of spatially selective cells necessarily produce an input to each CA1 cell that varies spatially as a realization of a random Gaussian process, due to the multivariate central limit theorem (see supplementary information A). Gaussian statistics arise robustly from random synaptic projections without the necessity to invoke highly specific assumptions on the structure of spatial responses in the input layer. These can range from regular tuning curves that tile the environment to heterogeneous and irregular tuning curves with a distance-dependent covariance function (see supplementary information A). Hence, the broad success of the model across data sets and statistics suggests that randomness, rather than specific design, governs the synaptic organization of inputs to CA1.

Recent experiments indicate that a dominant mode of plasticity in CA1 involves randomly occurring modifications of synaptic projections from CA3, triggered by inputs from the entorhinal cortex^42–44^. The randomness of the synaptic weights may be related to the cumulative effect of many such non-Hebbian plasticity events. Since all the measurements analyzed in this work were collected in highly familiar and uniform environments, it is possible that the stationarity of the underlying Gaussian processes is specific to such conditions. We speculate that more intricate features of the place cell code arise under richer behavioral conditions via activity-dependent plasticity mechanisms acting on top of a scaffold of random synaptic weights.

## Methods

### Mathematical Notation

Throughout the main text, methods section, and supplementary information, we use the mathematical notation defined below:

| Symbol | Description |
| --- | --- |
| $L$ | Length of one-dimensional (1D) environment |
| $D$ | Dimension of space |
| $L^D$ | Volume of environment for $D > 1$ |
| $x$ | Spatial coordinate |
| $\mathcal{GP}$ | Gaussian process |
| $h(x)$ | Gaussian process input to CA1 cell |
| $r(x - x')$ | Covariance function of $h(x)$ |
| $\theta$ | Normalized threshold |
| $\sigma$ | Correlation length |
| $f(x)$ | Firing rate of CA1 cells in the model |
| $\phi(\cdot)$ | Gaussian cumulative density function |
| $N$ | Field count per cell |
| $s$ | Field size |
| $\bar{s}$ | Field gap |
| $\chi$ | Euler characteristic |

### Model details

Place fields are modeled as rectified realizations of a stationary Gaussian process over *D* dimensional space (*x* ∈ ℝ^*D*^) with 0 mean and stationary covariance function *r*(*x* − *x*^′^),

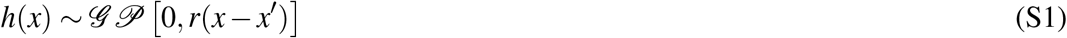

A sample *h*(*x*) from the Gaussian process can be thought of as input to a CA1 place cell. The input is subsequently thresholded at a value 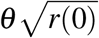 to generate the cell’s firing rate, which is proportional to *f* (*x*), defined as

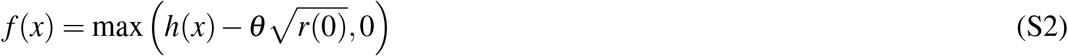

where *θ* is the normalized threshold, equal to the ratio between the threshold and the standard deviation of the process. The correlation length of the Gaussian process is defined as^26^:

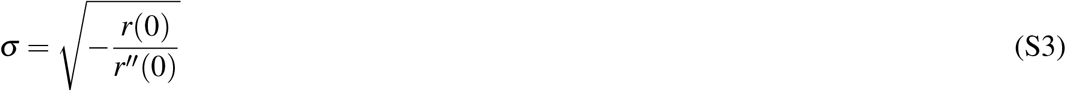

Thus, *σ* is equal to the ratio between the standard deviation of the process 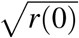 and the standard deviation of its derivative 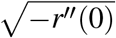^26^. For simulations of the Gaussian process, we used a transitionally invariant Gaussian correlation function.

### Data processing

Firing rate measurements from bats in 1d and 3d environments were first rectified at 0.5 hz for denoising. In the 3d case, firing rates also went through a median filtering using `scipy.ndimage` Python library with minimal (nearest neighbour) kernel of size 2 (place fields resolution = 10 cm on each axis) to eliminate salt-and-pepper noise while preserving field boundaries.

### Parameter inference in 1d environments

The two parameters *σ* and *θ* were inferred by matching the means of the field size and the field gap to those in the model via simulation in the data set from bats in the 200 m^6^ (Figs. 1C and 1D, Figs. 4A and 4B). In the dataset from bats in 6 m-long tunnel (Extended Data Fig. 5D), most cells had a single field and parameters were inferred by matching the mean field size and the mean fraction of the environment in which place cells were active. For the dataset from bats in the 130 m-long tunnel and dataset from rats in the 48 m-long maze, mean field size and mean field count were matched to infer the parameters^3,6^ (Figs. 4A, 4B, Extended Data Fig. 1). For rats in a 40 m-long virtual track^5^ (Fig. 1F), where only a histogram of field sizes was available, we directly fitted the parameter *β* of the approximate Rayleigh distribution (Eq. 1). Parameters matched through simulation yielded excellent agreement with the analytical formulas in the 1d case (Eqs. S21 and S22; see supplementary information).

### Parameter inference in 2d environments

For rats in the 2d environments^7^ (Figs. 1C-E and Figs. 4A,B), parameters were inferred by fitting the model to the mean field count and mean field size reported in^7^. When matching the mean count, we took into account the systematic exclusion of cells with zero fields from experimental observations by fitting the model to the distribution of field counts, conditioned on the existence of at least one field. For the Poisson distribution (valid in the high threshold regime), the mean ⟨*N*⟩_experiment_ of the conditioned distribution, which is matched to the experimental data, is then

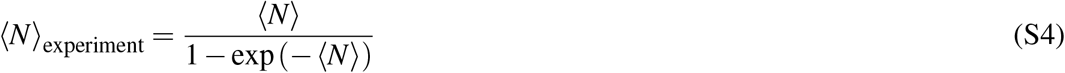

where ⟨*N*⟩ is the unconditioned mean. Analysis of the Euler Characteristic in the large 2D environment^7^ (Fig. 2H) was performed on a sample of firing rate maps from 20 cells shared with us by the authors of the manuscript. All other 2d results are based on statistics from the full data set, as reported in^7^.

### Parameter inference in 3d environment

For bats in a 3d environment^4^ (Figs. 2H-J and Fig. 4A) the parameters *σ* and *θ* were inferred by matching the mean field volume and the mean fraction of the volume in which place cells were active, based on the full rate maps.

### Fields in 1d

Fields were defined as connected components of non-zero activity, and gaps as connected components of zero activity (Figs. 1C 1D, 2B, 2E). When generating histograms based on the model (Figs. 1C, 2C, 2H-2J, 4C), errors bars were obtained by performing 10^4^ simulations of random fields using the model, each matching the number of cells to the data. Field size distributions from model simulations in 1d were compared against the best fit log-normal distribution by first generating a kernel density estimate of the field size distribution from the simulations (`scipy.stats.gaussian_kde`). The model has one less degree of freedom than the two-parameter log-normal distribution, since constraining the model parameters to also fit the gap size distribution reduces the parameter space to one dimension. Nevertheless, the log likelihood test indicated that the model fits the data better than the log-normal distribution (ΔLLR = −50.3 *<* 0). Differences in model complexity aware log likelihoods were measured using the Bayesian Information (BIC) and Akaike Information (AIC) criteria, yielding ΔBIC = −109.1 *<* 0 and ΔAIC = −104.5 *<* 0, which implies that the data fits the model better than log-normal.

### Kurtosis and skew

Kurtosis, quantifying the heaviness of distribution tails, was defined as

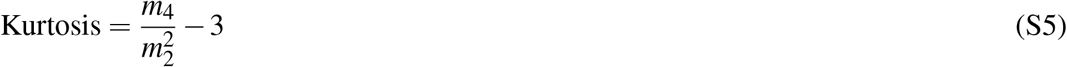

where *m*_4_ and *m*_2_ are the fourth and second moments of the distribution of log-field sizes, respectively. Skew, quantifying the asymmetry of the distribution was defined as

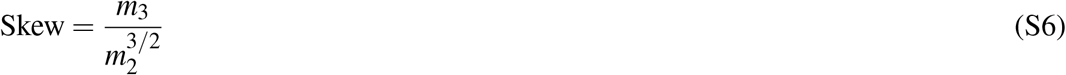

where *m*_3_ and *m*_2_ are the third and second moments of the log-field-size distribution, respectively. Both measures vanish for a log-normal distribution, since in that case the logarithrm of field sizes is normally distributed.

### Fields in 3d

Field statistics in 3D were measured from the firing rate data from^4^. To measure field volumes, connected components in the 3D volumetric fields were identified using the Python function `scipy.ndimage.label`, which identifies and labels connected regions in *n*-dimensional arrays. Fields with volume larger than 3 standard deviations from the mean were removed as outliers, as they are likely to result from merging of two or more fields. These large outliers were also removed when analysing the slices of the 3D fields (Figs. 2D, 2E, 2F).

### Number of local peaks within a field

Maxima in each field from 1d data were identified using the Python function `scipy.signal.find_peaks`, which finds local peaks in a 1D array. Incomplete fields at boundaries were removed. To filter out spurious maximas in the data arising from noisy assessment of the firing rate, we first applied a Gaussian smoothing with a standard deviation ranging from 0.5 m to 1 m in the case of bats flying in a 200 m tunnel, and a Gaussian smoothing with a standard deviation ranging from 0.3 m to 0.6 m in the case of bats flying in a 6 m tunnel. Local peak count distribution was calculated for the smoothed fields in each case and mean and standard deviation are reported. The error bars in the model represent the standard deviation across independent simulations (Fig 3B, 4C).

### Maximum firing rate vs. field width

Maximum firing rates in Figs. 3C, 3D were defined as the maximum rate observed in each field within its connected region. In the 1d data (Fig. 3C), fields were rectified by a small threshold (1 Hz) to eliminate spurious measurements arising from regions in which the firing rate measurement is unreliable (note that the Gaussian threshold crossing model is applicable for the rectified data). The power law coefficient extracted from the data was insensitive to the choice of the threshold above this value (Extended Data Fig. 7). The *y*-intercept of the linear fit on the log-log plot was matched to eliminate the effect of the linear gain factor.

### Boundary slopes

Field boundary slopes were extracted by fitting a line between the two consecutive samples of the firing rate (separated by 0.1 m) at the boundary of each rectified field, and measuring its slope (Fig 3E).

### Euler characteristics

The Euler characteristic (EC) is a topological invariant that can be expressed as the alternating sum of Betti numbers. The 0-th order Betti number is equal to the number of connected components in a topological space, and the *n*-th order Betti number is equal to the number of *n*-dimensional holes. The EC *χ* of a Gaussian process in *D* dimensions has a universal form, insensitive to the structure of the underlying covariance function (under mild regularity requirements,^26^):

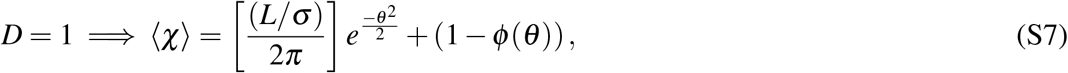

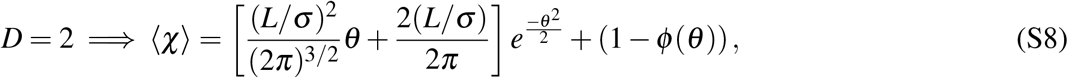

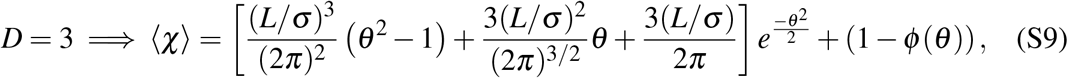

where *ϕ* is the Gaussian CDF function. Empirical Euler characteristics (EC) were measured using the `Gudhi` python library by taking the alternating sum of Betti numbers. The mean maximum firing rate was normalized in the data to match that of the model to determine the linear gain value before calculating the EC.

## End notes

## Acknowledgments

We thank Tamir Eliav, Shir Maimon, and Nachum Ulanovsky for sharing with us full data sets comprising rate maps of CA1 cells in bats from^6^ and^4^, and for providing helpful information on these data sets. We thank Nachum Ulanovsky for helpful discussions and for comments on the manuscript. We thank Jean-Marc Fellous for sharing with us a sample data set of rate maps from 20 CA1 cells in rats from^7^.

## Extended Data

**Exdented Data Fig. 1.**
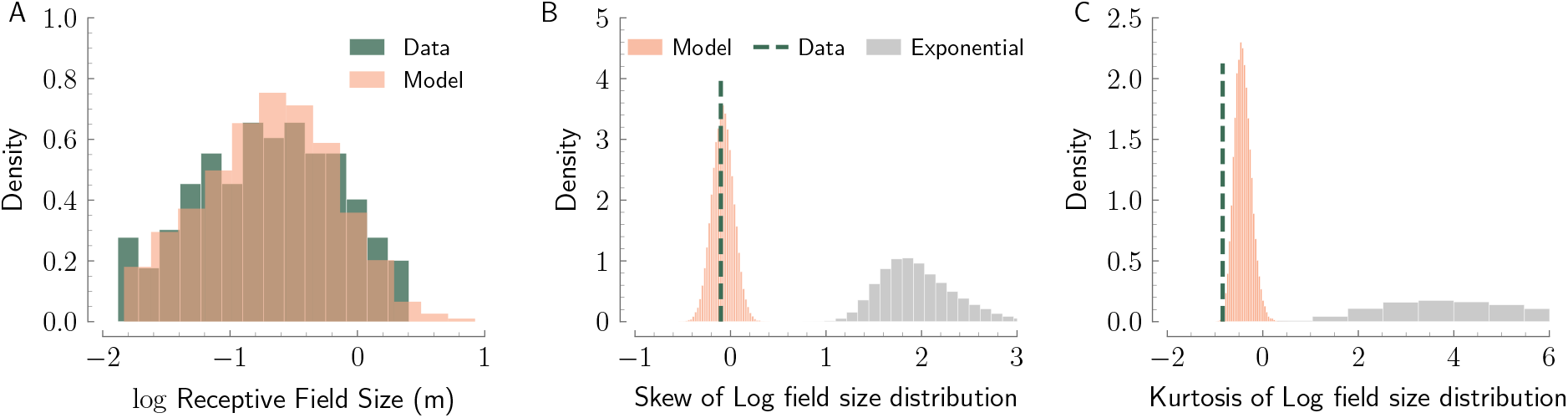
**a**, Distribution of (log) field sizes measured in rats navigating a 48 m-long 1d maze^3^ along with distribution arising from the model (*σ* = 0.34, *θ* = 1.8), fitted to match mean field size and count observed in the data. Fields in data larger than three times the standard deviation were removed as outliers. The empirical distribution matches the model better than the heuristic exponential distribution proposed in^45^ (likelihood test: Δlog Likelihood = − 83.1 *<* 0). **b-c**, Skew/Kurtosis distribution of log field size obtained from multiple simulations of the model (orange) and the heuristic exponential distribution (gray) compared against that extracted from the experimental data (vertical green dashed line). The empirical fields were required to be larger than 15 cm in^3^. We applied the same cutoff on the fields from to the model to match the distribution obtained from the data. Note that this affects the distribution of skew and kurtosis.

**Exdented Data Fig. 2.**
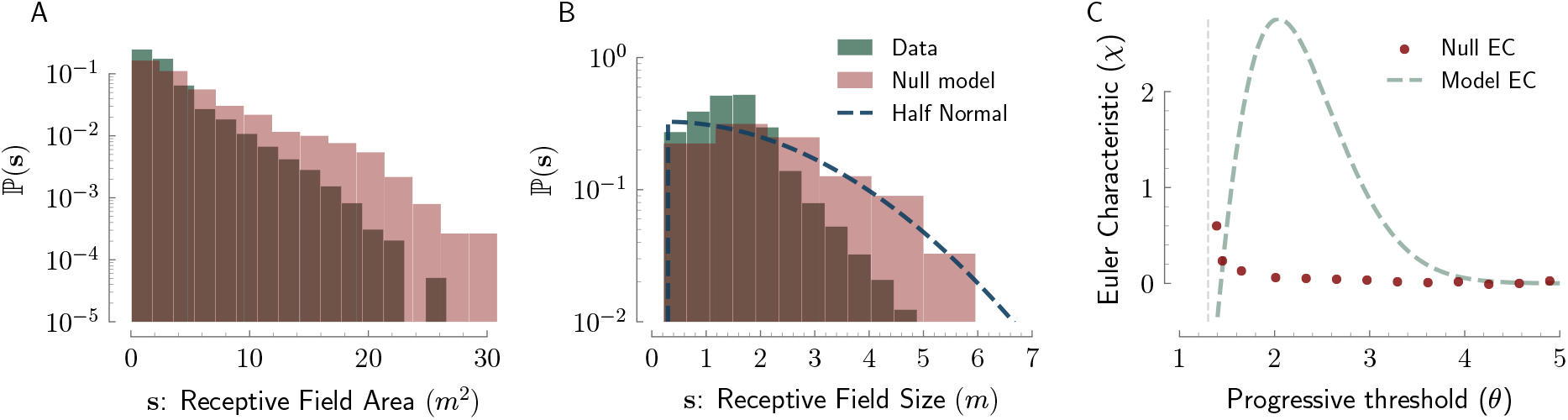
Alternative model for comparison against the thresholded Gaussian process model in 3d. Firing fields in the alternative model are spherical, while the joint distribution of field volumes and maximum-firing rates exactly matches the empirical distribution. More precisely, for each field in the data, we created a corresponding field in the model with spherical support and matching volume. These fields are not allowed to overlap but are otherwise randomly placed since the measurement of field sizes in the slices and the expected Euler characteristics are independent of the field placements. The firing rate profile for each of these fields was modeled as a quadratic function with amplitude exactly matching that of the field in data. **a**, Discrepancy between areas of 2d slices through 3d fields in data and in the alternative model (logarithmic vertical scale). **b**, Discrepancy between length of 1d slices through 3d fields in data and in the alternative model (logarithmic vertical scale). In the null model, the radius of each spherical field is distributed as Rayleigh distribution, which implies that the length of active portions of 1d slices follows a half-normal distribution. **c**, Comparison between Euler characteristics of the thresholded Gaussian process model (matching the empirical data, Fig 3G), and Euler characteristics arising from the alternative model.

**Exdented Data Fig. 3.**
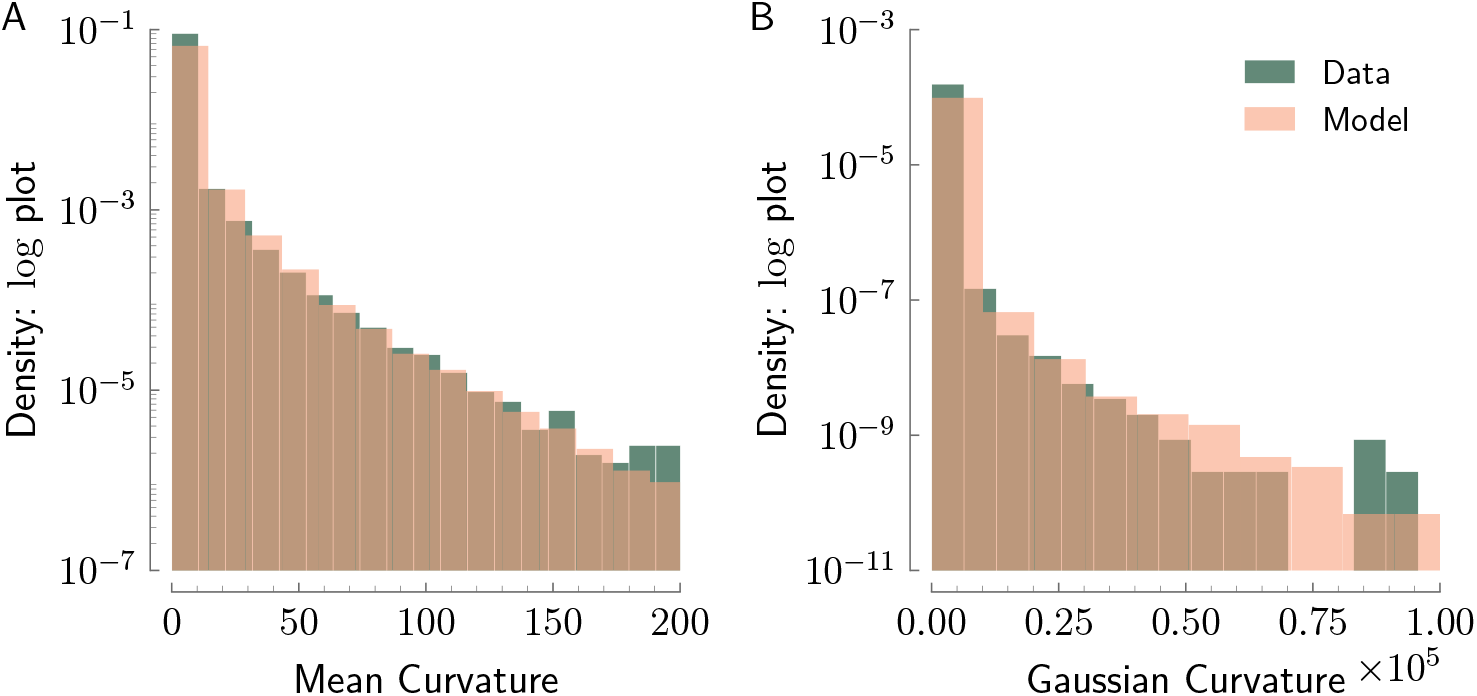
Distribution of mean curvature (**a**) and Gaussian curvature (**b**) of 3d place field boundaries in data and model (logarthmic vertical scale). The boundary surface for each field was first identified and represented as a two-dimensional array *Z*, where each point (*x, y, z*) of the surface was mapped to position (*x, y*) in the array containing the *z*-value of the surface points. The mean and Gaussian curvatures of the isosurface representing the boundaries of the 3D fields were then calculated using the following operators^46^: Gaussian curvature (*K*) is defined as:

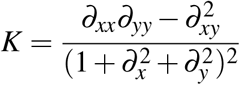

and mean curvature (*H*) is defined as

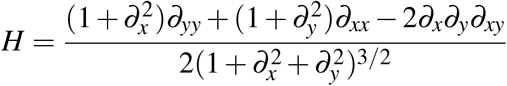

where *∂*_*x*_, *∂*_*y*_, *∂*_*xx*_, *∂*_*yy*_, and *∂*_*xy*_ are the first and second-order partial derivatives of the surface represented by array *Z* with respect to *x* and *y*. The partial derivatives were computed using the `numpy.gradient` function, which calculates the numerical gradients of the input array.

**Exdented Data Fig. 4.**
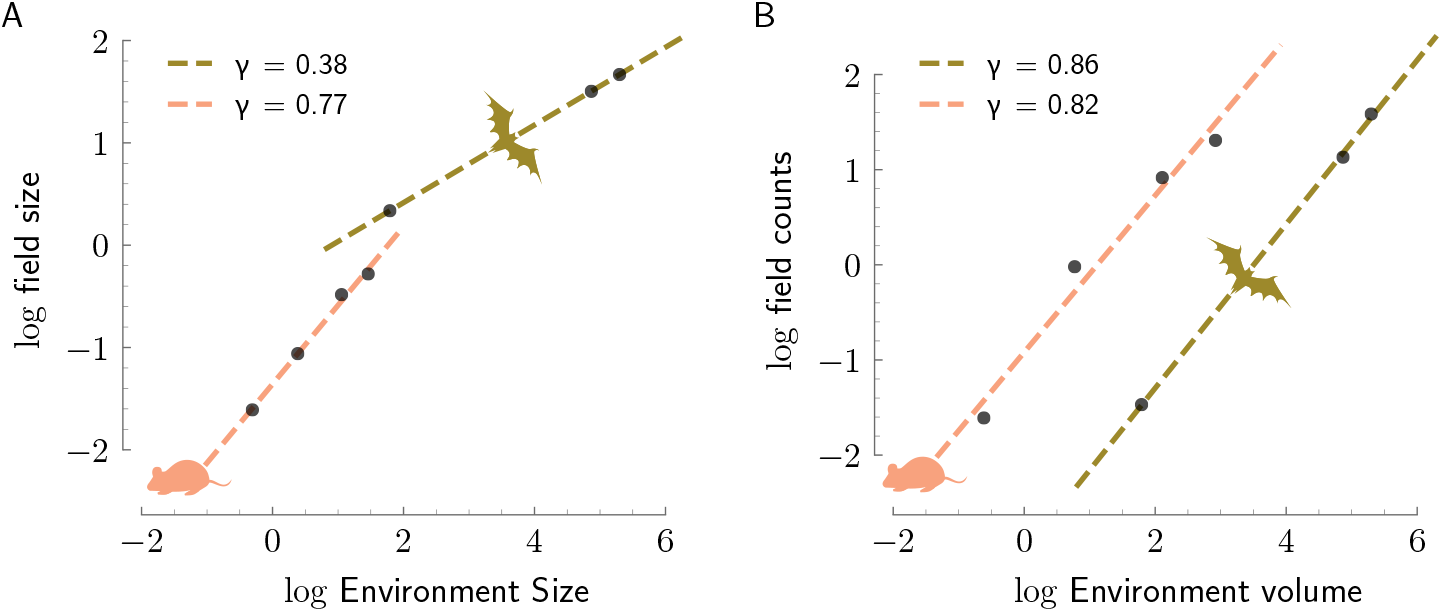
**a**, Linear scaling of logarithm of linear field sizes (in m) with the logarithm of the linear environment size (in m) in 1D (bats)^6^ and 2D (rats)^7^. **b**, Linear scaling of logarithm of field counts with the logarithm of the linear environment volume (length in 1d and area in 2d) in 1d (bats)^6^ and 2d (rats)^7^. In both panels, *γ* is the slope of the best linear fit to the data in logarithmic scales.

**Exdented Data Fig. 5.**
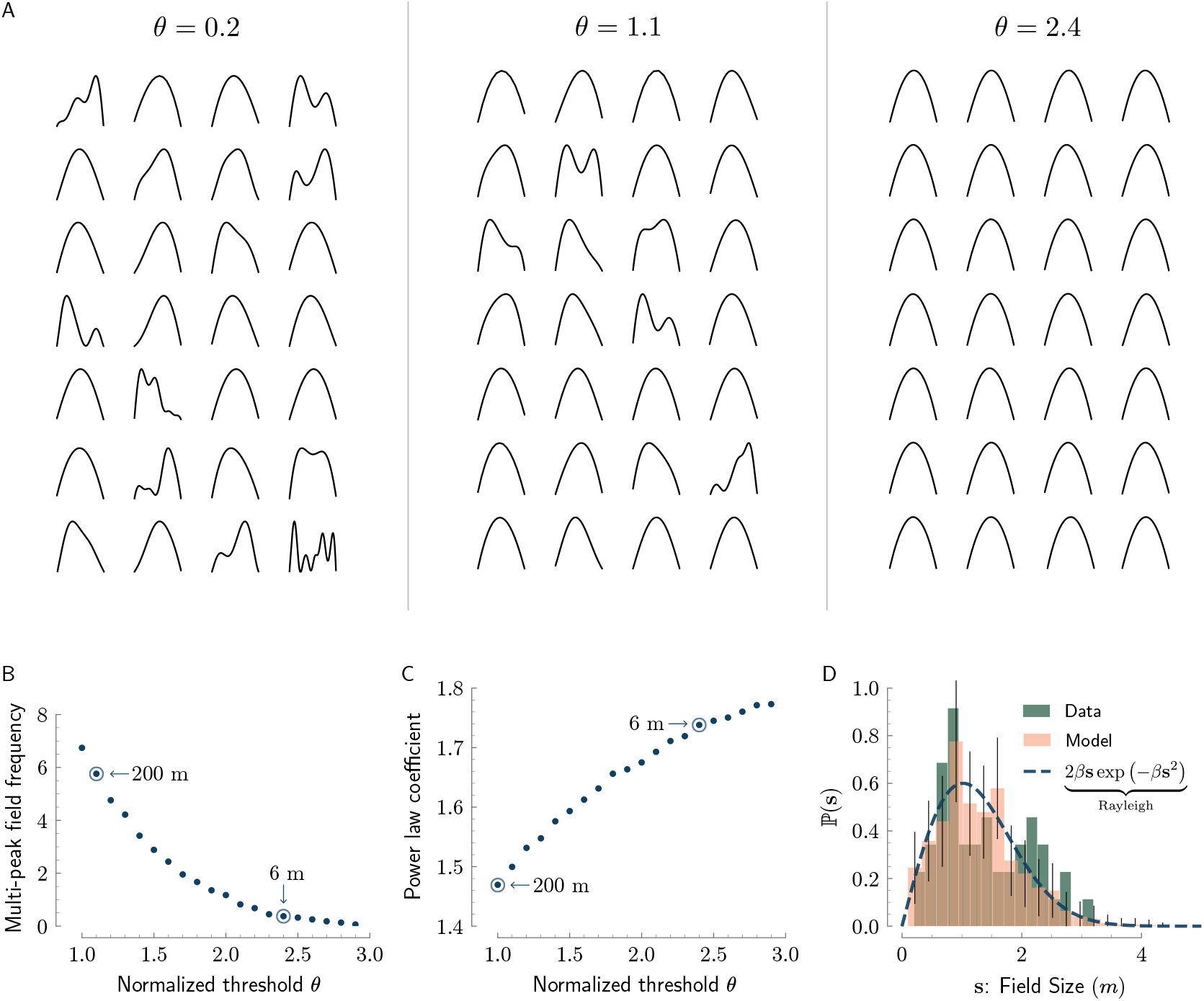
**a**, Visualization of sample field shapes generated by the model ranging from low to high threshold regime (left to right. The intermediate value of *θ* = 1.1 is the normalized threshold inferred from recordings in the 200 m-long tunnel). In all cases, fields were randomly selected. **b**, Predicted percentage of cells with more than one local peak, as a function of the normalized threshold. **c**, Predicted logarithmic slope of field width vs. max. firing rate, as a function of the normalized threshold. It can be shown that in the limit of very high threshold, fields are parabolic and the logarithmic slope approaches 2^26^. **d**, While the theory predicts that field shapes are more stereotyped for higher thresholds as described in panels A,B, it predicts that field widths follow the Rayleigh distribution for large thresholds. Therefore, the variability in field widths relative to their mean is insensitive to the threshold. This is corroborated by the empirical distribution of field widths from bats flying in a 6 m-long tunnel. Note that the measurement is noisy due to the small number of cells (40 recorded cells; peak count in the figure is 8; the distributions have no significant difference; K-S test: p-value=0.6, k-s stat=0.1).

**Exdented Data Fig. 6.**
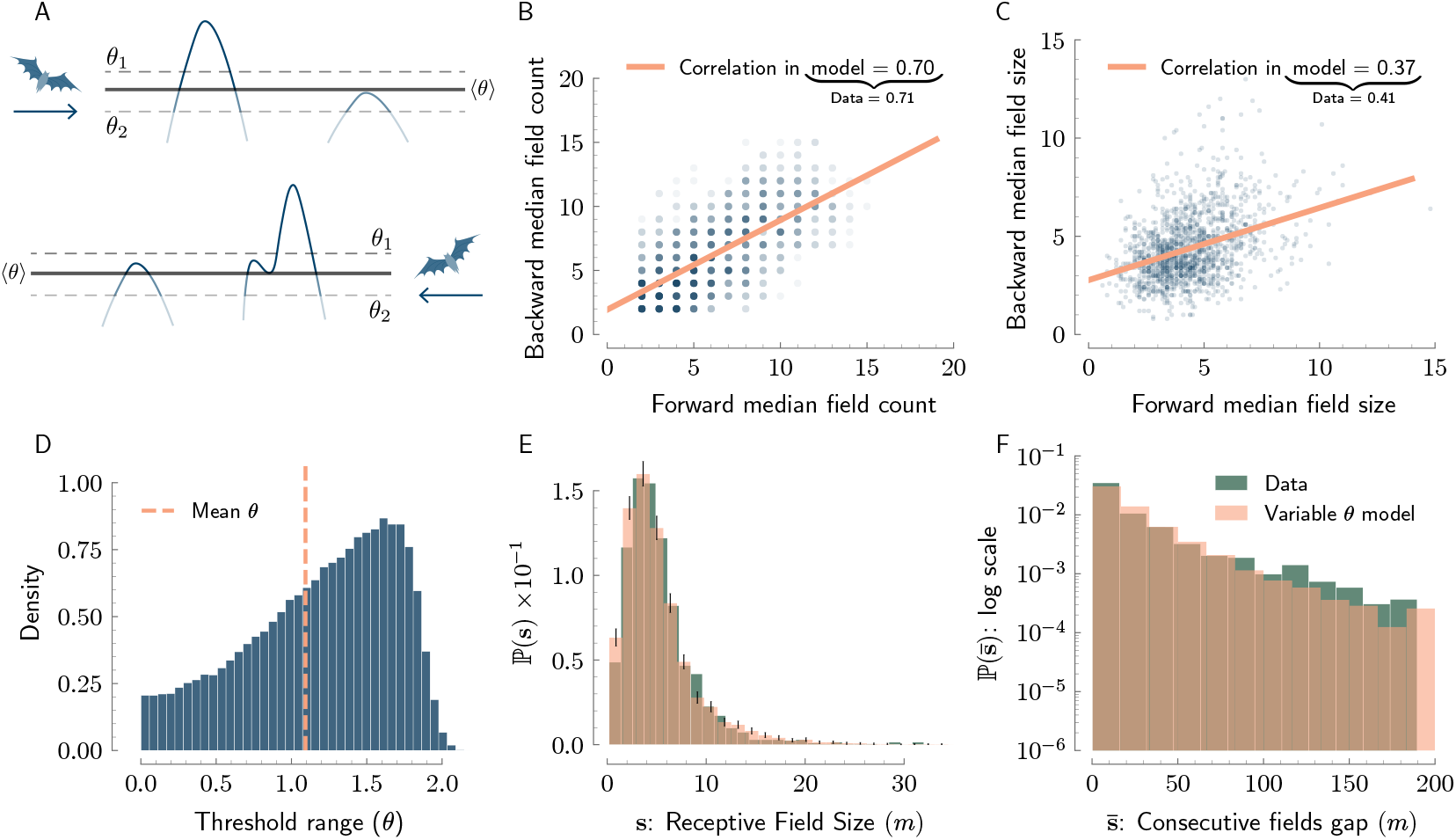
**a**, Illustration of how variability in the normalized threshold across cells leads to correlated field counts and field sizes in the two flight directions along the 1d tunnel. Neurons with lower threshold will have more place fields on average, when the bat flies along both directions of the tunnel. Likewise, neurons with lower threshold will have larger field size on average, when the bat flies along both directions of the tunnel. **b**, Scatter plot (blue) and correlation of field counts obtained from the model in two different samples (representing forward and backwards directions) of fields with fixed noisy threshold. With a distribution of thresholds, the model matches the empirical correlation between field counts in bats flying in the forward and backward directions (parameters chosen as explained in D). **c**, Scatter plot (blue) and correlation of median field size per cell in two different samples of fields with fixed noisy threshold. With a distribution of thresholds, the model matches the empirical correlation between field sizes in the two flying directions, while simultaneously matching the correlation in field counts (parameters chosen as explained in D). **d**, The distribution of *θ* in the extended model with variability in the threshold across cells. *θ* is sampled from a skew-normal distribution conditioned to be positive, with mean matching the single inferred *θ* in case of bats in 1d tunnel. The skew normal distribution has two remaining parameters after fixing the mean, which were set to match the correlation between median field count and median field size as reported in^6^. **e**, Field size distribution under the variable threshold model matches the distribution of field sizes in the data. **f**, Field gap distribution in the model matches that of the data and further explains (without additional fitting) the slightly heavier tail in the empirical distribution in comparison with the fit to an exponential distribution (compare with Fig. 1D).

**Exdented Data Fig. 7.**
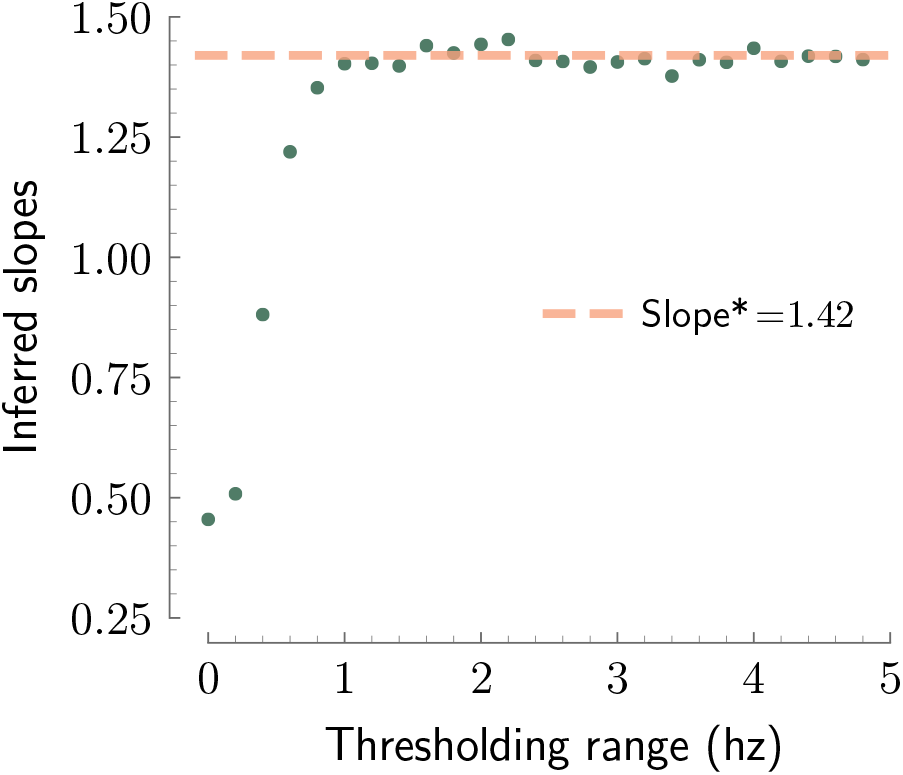
Slope of the linear fit between log field size and log maximum firing rate, evaluated for the data from bats flying in the 200 m tunnel using a range of rectification parameters. For rectification parameters above 1 Hz the slope rapidly stabilizes on a consistent value.

## Supplementary Information

### A. Derivation of Gaussian process input to CA1 cells

Here we illustrate how random summation of presynaptic inputs into CA1 cells leads to Gaussian statistics, and elucidate how the spatial correlation function of the Gaussian process is determined by the input statistics. We assume, for simplicity, completely independent weights, yet Gaussian statistics can arise also under broader conditions, in which there is partial correlation between groups of summed variables that decays sufficiently rapidly across a functional axis within the input population^47^.

Consider a population *i* = 1 … *N* of spatially selective neurons presynpatic to a CA1 place cell with bounded spatial response function *u*_*i*_(*x*) for *x* ∈ ℝ^*D*^ in *D* dimensions. For simplicity we consider these response functions to have zero mean, whereas in a more realistic setting *u*_*i*_ can stand for the deviation of the firing rate from its mean, and approximate balancing of the mean (which is interchangeable also with the choice of threshold in our model) may be obtained by other mechanisms, such as mutual lateral inhibition or homeostatic regulation of excitatory and inhibitory inputs.

We consider first, for simplicity, a scenario in which all input neurons have the same tuning curve up to translation, with preferred firing locations *x*_*i*_ that uniformly and densely tile the space:

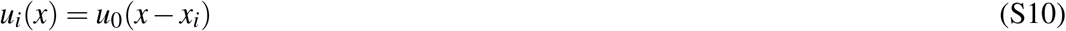

The input to the CA1 cell, *h*(*x*) is a weighted linear sum of the activity of the presynpatic cell outputs:

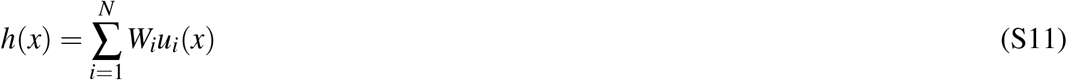

where we assume that the weights *W*_*i*_ are drawn independently from a random distribution *p*(*W*) with zero mean, and a finite second moment. Using the central limit theorem for multidimensional stimuli^48,49^, it is straightforward to see that for any discrete set of locations *x*_*α*_, *h*(*x*_*i*_) are jointly distributed as multivariate normal in the large *N* limit. Hence, *h*(*x*) is a Gaussian process in this limit.

The covariance function of the Gaussian process (which uniquely determines all of its properties) must be translationally invariant, due to our assumption of uniform tiling of inputs in the input layer. Indeed,

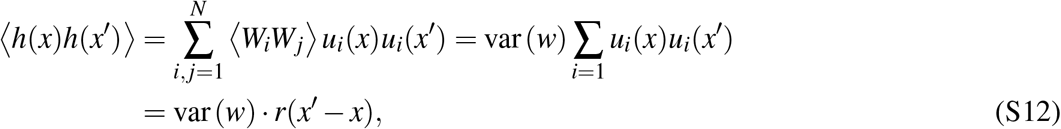

where the angular brackets represent an average over realizations of the random synaptic weights, and *r* is the spatial covariance of the neural population response. Here, *r* is proportional to the auto-correlation of the single neuron tuning curve:

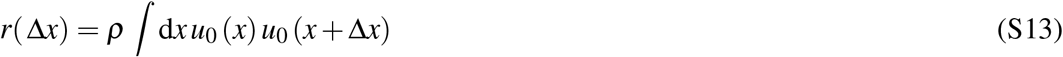

where *ρ* is the spatial density of receptive field centers. Specifically, if individual tuning curves are shaped as Gaussian firing fields, *r*(Δ*x*) is a Gaussian with a width equal to twice the width of the individual firing fields.

#### Heterogeneous tuning curves

Similar results hold for heterogeneous tuning curves. We assume that the tuning curves of individual neurons are sampled from a distribution of shapes, parameterized by a random variable *α* which is independently chosen for each neuron (*α* can be scalar or multidimensional, discrete or continuous). For each neuron *i*,

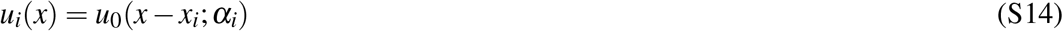

where *x*_*i*_ is drawn from a a uniform distribution in space and *α*_*i*_ is the shape parameter of neuron *i*. Under these assumptions, *h*(*x*) is a sum over many independent random functions, which becomes distributed in the large *N* limit as a Gaussian process due to the central limit theory. A similar derivation as above yields

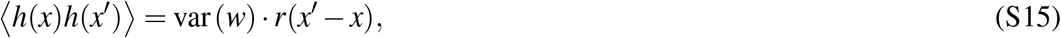

where *r*(Δ*x*) is proportional to the auto-correlation of the tuning curve, averaged over the distribution of receptive field shapes:

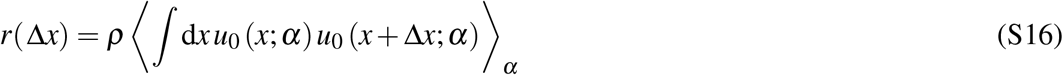

where ⟨⟩_*α*_ represents an average over realizations of the shape parameter *α*. Qualitatively, in both scenarios discussed above, the central limit theory becomes valid when each CA1 neuron receives, within its place fields, active presynaptic inputs from many spatially selective cells.

### B. Field statistics

An extensive body of mathematical results exist for the statistics of excursion sets of Gaussian processes. In this section we explain how a small subset of the results are obtained. Full derivations of all the results are found in^26^.

#### Field statistics in 1D

Field counts in 1d can be exactly derived using the Kac-Rice formula^32^,

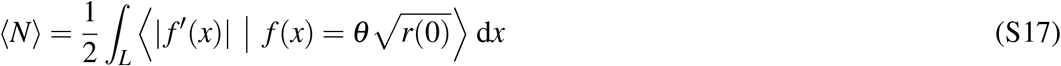

where *f* (*x*) is a random process with a well defined derivative. Specifically, in our model *f* (*x*) is a 0 mean, stationary Gaussian process, and consequently *f* ^′^(*x*) is a 0 mean stationary Gaussian process as well. To calculate the integrand, we first compute the covariance matrix associated with the joint distribution of *f* (*x*) and *f* ^′^(*x*),

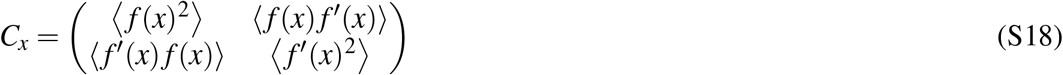

Using stationarity, we have:

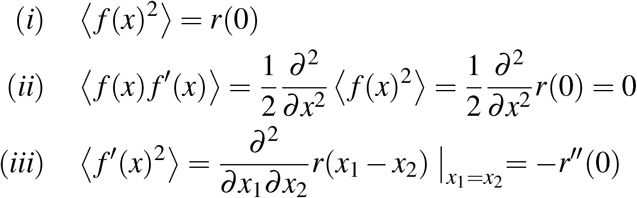

Thus,

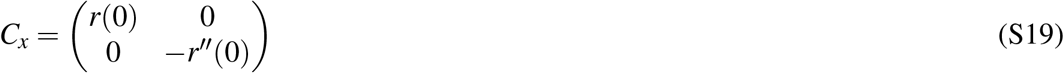

Using (S19) we can calculate the expectation inside the integral,

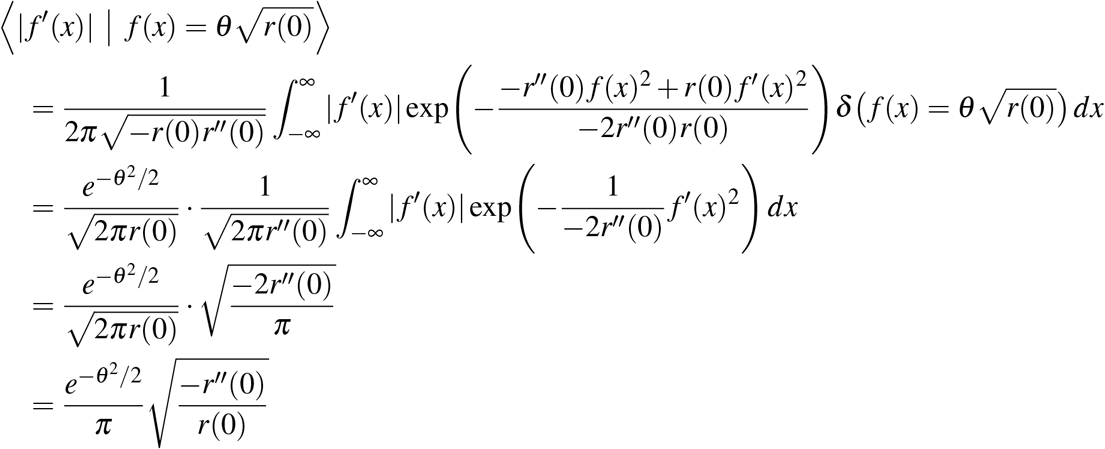

Therefore, the final result for ⟨*N*⟩ is:

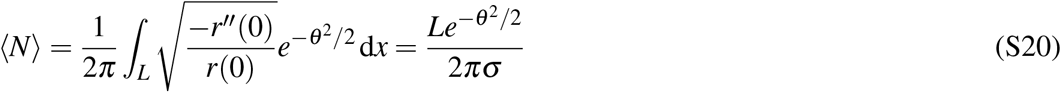

Next, assuming that the environment is large compared to the correlation length, the portion of the environment in which a cell is active is *L* (1 − *ϕ* (*θ*)). This, together with the mean field count can be used to calculate the mean field size *s* and the mean gap between the fields 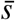as:

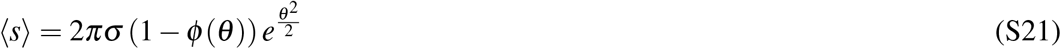

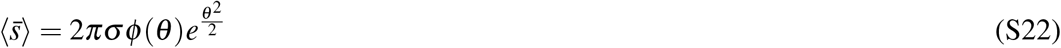

Simulations were in excellent agreement with all these expressions.

#### Field statistics in higher dimensions

In higher dimensions, precise analytical expressions are not available for the mean field count and size. We relied on simulations to determine the parameters *σ* and *θ* that match the mean size and the mean field count, as described in Methods. A qualitative understanding of these relationships can be obtained using an approximation which is valid for high thresholds, obtained by retaining the leading order term in the expression for the Euler characteristics (see^26^ for more details on this approximation):

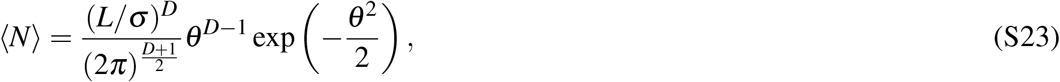

where *L*^*D*^ is the volume of the environment.

The mean field size is then approximated by

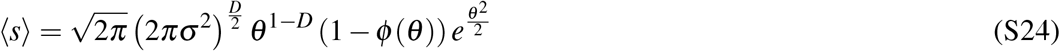

#### Distributions in the high threshold regime

As described in the main text, distributions related to the field arrangements acquire a universal form in the high threshold regime. The field count is Poisson, in all dimensions, with a mean given by Eq. S23. This is equivalent to the fact that the field gaps in 1d are exponentially distributed. Similarly we can approximate the full field-size distribution as in the main text (Eq. 1) by:

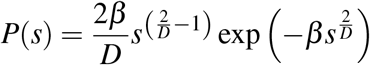

Furthermore, we can relate *β* to the parameters *σ* and *θ* of the model through the analytic forms for the mean field size (Eqs. S22 and S24). To that end, first, observe that *s*^2*/D*^ is exponentially distributed with an expectation ⟨*s*^2*/D*^ = 1*/β ⟩*, implying that ⟨*s*⟩ = Γ(*D/*2 + 1)*β* ^−*D/*2^ using the moment generating function of the exponential distribution. Therefore:

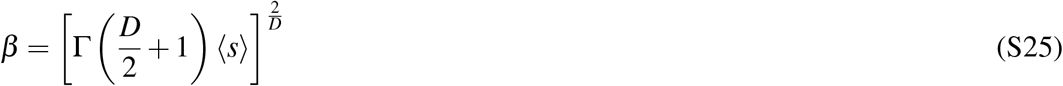

